# VEF: a Variant Filtering tool based on Ensemble methods

**DOI:** 10.1101/540286

**Authors:** Chuanyi Zhang, Idoia Ochoa

## Abstract

**Motivation:** Variant discovery is crucial in medical and clinical research, especially in the setting of personalized medicine. As such, precision in variant identification is paramount. However, variants identified by current genomic analysis pipelines contain many false positives (i.e., incorrectly called variants). These can be potentially eliminated by applying state-of-the-art filtering tools, such as the Variant Quality Score Recalibration (VQSR) or the Hard Filtering (HF), both proposed by GATK. However, these methods are very user-dependent and fail to run in some cases. We propose VEF, a variant filtering tool based on ensemble methods that overcomes the main drawbacks of VQSR and the HF. Contrary to these methods, we treat filtering as a supervised learning problem. This is possible by using for training variant call data for which the set of “true” variants is known, i.e., a *gold standard* exists. Hence, we can classify each variant in the training VCF file as true or false using the gold standard, and further use the annotations of each variant as features for the classification problem. Once trained, VEF can be directly applied to filter the variants contained in a given VCF file. Analysis of several ensemble methods revealed random forest as offering the best performance, and hence VEF uses a random forest for the classification task.

**Results:** After training VEF on a Whole Genome Sequencing (WGS) Human dataset of sample *NA12878*, we tested its performance on a WGS Human dataset of sample *NA24385*. For these two samples, the set of high-confident variants has been produced and made available. Results show that the proposed filtering tool VEF consistently outperforms VQSR and HF. In addition, we show that VEF generalizes well even when some features have missing values, and when the training and testing datasets differ either in coverage or in the sequencing machine that was used to generate the data. Finally, since the training needs to be performed only once, there is a significant saving in running time when compared to VQSR (50 minutes versus 4 minutes approximately for filtering the SNPs of WGS Human sample NA24385). Code and scripts available at: github.com/ChuanyiZ/vef.

## 1 Introduction

Next Generation Sequencing (NGS) technology is becoming extremely cost-effective, making personalized genomic medical research a reality. Due to the high similarities across genomes of the same species, most of the medical and clinical analyses start with variant calling, i.e., the identification of variants (differences) of the sequenced genome with that of a reference genome. In particular, variant identification from Whole Genome Sequencing (WGS) data is increasingly being used to diagnose, gain biological insight to, and design treatments in the clinical setting. Thus, precision in variant identification is paramount.

The process of identifying the variants starts with the raw data produced by NGS technologies, which are stored in the FASTQ format. The raw data consist of millions of short fragments of nucleotides from the sequenced genome, called reads, each accompanied by a sequence of quality scores and an identifier. These fragments are then generally aligned to a reference sequence of the same species of the genome that was sequenced, generating an aligned SAM/BAM file [Li *et al.*, 2009]. This file is then fed as input to the variant caller, which generates the set of variants believed to be present in the sequenced sample. These variants are generally given by Single Nucleotide Polymorphisms (SNPs), and insertions and deletions, the latter referred to as INDELs. The called variants are currently stored in VCF (Variants Call Format) files [Danecek *et al.*, 2011], which also contain significant metadata (in the form of annotations) related to the process of calling these variants (see Figure 1 for an example). More details on the format are given in Section 2.

**Figure 1:**
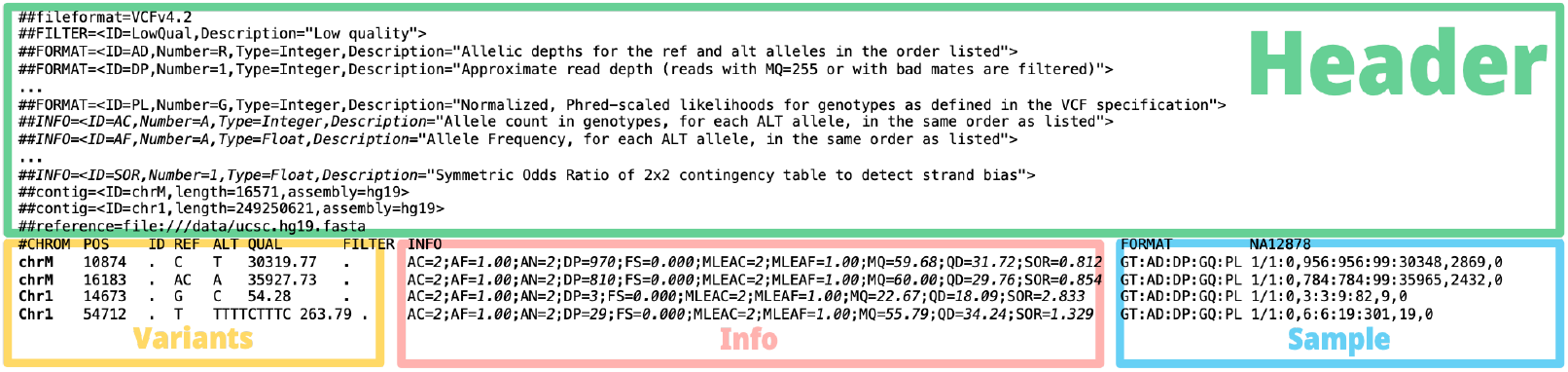
Example of a VCF file, extracted from Human sample *NA12878*, sequenced by 10 × Genomics and processed with GATK v3.8. In this case, the INFO field contains 10 annotations, given by AC, AF, AN, DP, FS, MLEAC, MLEAF, MQ, QD, and SOR.

Unfortunately, variants identified by current genomic analysis pipelines contain many incorreclty called variants (i.e., false positives) and miss several true ones. For example, the variant datasets analyzed in [Ochoa *et al.*, 2016] contain on average more than 25% false positives. These errors may come from the sample preparation, the errors introduced during the sequencing process, and the heuristics of the methods used in the analysis pipeline (including the alignment and variant calling tools).

To further reduce the number of incorrectly called variants in the generated VCF files, the Genome Analysis Toolkit (GATK) [DePristo *et al.*, 2011] provides the filtering tools VQSR (Variant Quality Score Recalibration) and Hard Filtering (HF), the latter recommended in cases where VQSR is not suitable. Due to the different characteristics of SNPs and INDELs, both VQSR and the HF are applied to each of them separately. These methods are considered the state-of-the-art, and hence next we explain them in more detail.

### VQSR

In brief, VQSR first selects the subset of variants in the VCF file that are known to exist on highly validated variant resources as the training set (e.g., for Human data, Omni, HapMap [Consortium *et al.*, 2003], and 1000 Genomes [Consortium *et al.*, 2015]). The rationale is that these variants are probably correctly called. VQSR then trains a Gaussian mixture model (GMM) in these variants, using a set of user-selected annotations as features (note that for each variant in a VCF file a set of annotations is provided). Once the GMM is trained, the model is applied to all variants in the VCF file, and a recalibrated quality score is computed for each of them. This recalibrated quality score, also known as variant quality score log-odds (VQSLOD), indicates the probability that the variant was generated by the fitted GMM, or in other words, the probability that the variant is true. As the last step, the variants with a low VQSLOD value are filtered out. More specifically, the user specifies a tranche sensitivity threshold, expressed as a percentage (e.g., 99.9%), that VQSR uses to determine the VQSLOD value above which 99.9% (in this specific example) of the variants in the training set are included. This value is then used to filter the variants with a smaller VQSLOD.

VQSR has several limitations, including a heavy dependence on user-selected parameters. For example, the number of Gaussians in the GMM, the specific annotations to be used for training, or the value of the tranche sensitivity threshold, the latter directly affecting the sensitivity of VQSR. In addition, as VQSR needs to train on the target VCF file, it does not work on small files containing less than 1000 variants. Similarly, VQSR cannot be applied in cases for which true/training sets are not available. In these cases, the HF approach is recommended instead.

### HF

The HF approach is the most basic and traditional way of filtering variants. Given a set of user selected annotations and corresponding thresholds, the hard filtering approach will filter out those variants that do not satisfy the given constraints, i.e., those with annotation values above or below (depending on the annotation) the set thresholds. The main problem with this approach is that it is very limiting, as the value of a single annotation may filter out a variant, even if all other annotations have values within the given thresholds. For example, the HF may throw out all variants with DP (read depth) annotation below the set threshold, independently of the value of the other annotations. In addition, its performance depends heavily on the user, who may not have the right or necessarily knowledge to decide on the most suitable set of annotations and thresholds.

With the goal of overcoming the main limitations of these methods, here we propose VEF, a new variant filtering tool. Unlike VQSR and HF, VEF approaches the filtering of variants as a supervised learning problem, and it is based on ensemble methods. In particular, VEF trains a random forest on a set of variants for which the information regarding which variant is correct and which one is not is known and available (i.e., for which the *ground truth* or *gold standard* exists). This is the case for example of Human sample *NA12878*, for which a high-confidence variant call set has been provided by the National Institute of Standards and Technology’s (NIST) Genome In A Bottle (GIAB) consortium [Zook *et al.*, 2018]. We make the assumption that VCF files generated by the same analysis pipeline share the same feature space and characteristics. Hence, under this assumption, once the model is trained, VEF can be directly applied to any VCF file generated by the same pipeline. As such, VEF does not need any pre-calibration step by the user, and it can be applied on arbitrary small VCF files. As an added advantage, since the model needs to be trained only once, significant savings in running time are achieved with respect to VQSR. We further show on real data that VEF outperforms VQSR and HF in classification accuracy, including cases in which VEF is trained and tested on VCF files generated from raw data of different coverages or sequenced with different sequencing machines, and when tested on VCF files containing features with missing values.

It should be noticed, however, that VEF is designed for filtering variants from single non-cancerous samples, and hence for large cohorts or cancerous samples, other methods may be preferred.

## 2 Methods

VCF files consist of a header section followed by a record section. The header section contains information related to the VCF file itself, such as the version and the commands used to generate it, among others. The record section contains the called variants, including both SNPs and INDELs. Information for each variant is specified in a separate line, and contains the location of the variant (chromosome and position), its ID, the reference base(s) and alternative base(s), a quality score, filter information, annotations, and sample information. Annotations are generally stored in the “INFO” field, with names and types specified in the header. Figure 1 shows a snapshot of a VCF file. The version of VCF used in this work is v4.2 (the latest), documented at https://samtools.github.io/hts-specs/VCFv4.2.pdf.

Given a VCF file containing a set of called variants, the goal of the variant filtering tools is to filter out the incorrectly called variants while retaining the true ones. This decision is generally made as a function of the annotations (features) associated to each variant. Currently available filtering tools differ in how this decision is made. Nevertheless, all previously proposed classifiers, such as VQSR and HF, are based on unsupervised or semi-supervised learning methods. However, with the release of high-confident variant call sets of various Human samples, such as those released by the NIST’s GIAB consortium [Zook *et al.*, 2018], it is now possible to utilize supervised learning methods for variant filtering/classification. The rationale is that VCF files generated with the same analysis pipeline will exhibit a similar probability distribution of the annotations of incorrectly and correctly called variants (note that in a way, the HF also makes this assumption). Hence, under this assumption, the model needs to be trained only once for each considered pipeline, and then it can be directly applied to any VCF file generated by the same pipeline.

One of the main advantages of using supervised learning include being able to apply the filtering tool to any VCF file, regardless of its size, without any input from the user. In addition, the elimination of the training phase for every new VCF file that needs to be filtered can significantly speed up the analysis.

To benefit from these advantages, the proposed variant classifier VEF is based on supervised learning. In this sense, the filtering of variants could be done by using any supervised learning binary classification technique. However, we restrict our attention to ensemble learning methods since they are suitable for the problem at hand, as explained next. Ensemble methods compromise a series of base learners, usually Decision Trees (DTs), whose predictions are aggregated to produce the final prediction. A decision tree is a classification method represented by a binary tree structure. Each internal node of the tree uses a rule to split the dataset into two subsets, represented by the two child nodes of that node. The rule is usually defined as the value of a feature being above or below a given threshold. By iteratively testing on the rules, the feature space is cut into several regions. Each region, corresponding to a leaf node in the tree, is associated with a class. For the problem at hand, the classes correspond to the variant being correct or not. Decision trees have low bias and tend to overfit the data easily. However, by combining the prediction of each DT using majority or weighted voting to make the final prediction, ensemble methods can have both low bias and smaller generalization error, which is suitable for classification [James *et al.*, 2013]. The final decision of ensemble methods is therefore made by choosing the class with the highest weighted probability. For binary classification, since the probability of the two classes sums up to one, the default threshold (or cut-off value) is 0.5. Another advantage of using decision trees as base learners is that data normalization is not required, which is useful when normalization itself introduces bias.

In this work we consider the most common ensemble methods used in practice: Random Forest (RF) [Breiman, 2001], AdaBoost [Freund and Schapire, 1997], and Gradient Boosting Decision Tree (GBDT) [Friedman, 2001]. The main difference between these methods is the manner in which they train the base learners (decision trees). RF independently draws training data for each base learner and the final result is computed as the arithmetic average of the output of all decision trees. Both AdaBoost and GBDT add new base learners sequentially and make the final decision based on a weighted vote. AdaBoost and GBDT differ in the way they build new base learners. AdaBoost assigns weights to every sample point, and subsequently increases the weights of the miss-classified samples. Samples are selected based on the assigned weight, and hence the rationale is that subsequent added classifiers will focus more on the “difficult” samples. GBDT uses gradient descent instead. It computes the gradient of the loss function and builds a new learner along that direction with a weight (step size) which minimizes the loss. The general structure of ensemble methods is depicted in Figure 2.

**Figure 2:**
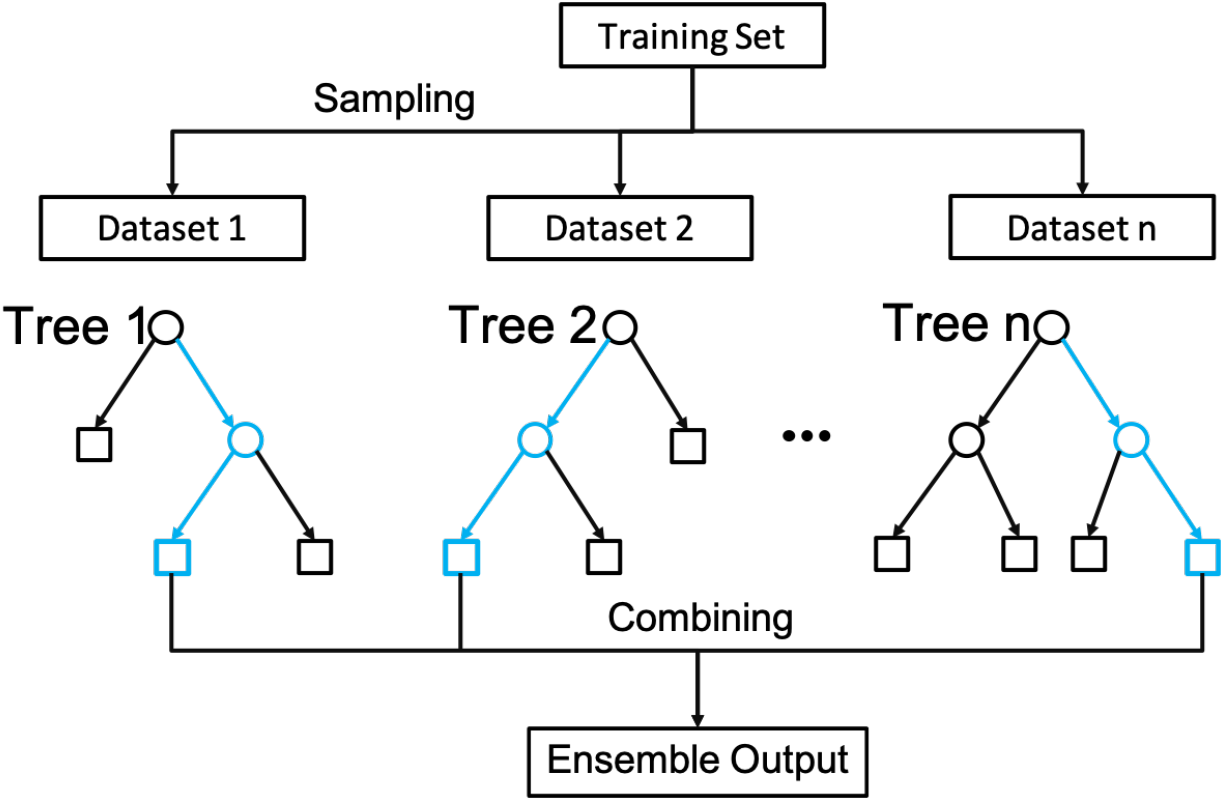
General structure of ensemble methods using decision trees as base learners. To generate the final prediction, RF uses majority voting, while AdaBoost and GBDT use weighted voting.

Among the considered ensemble methods, RF with 150 base learners was selected for the proposed filtering tool VEF, as it showed the best classification performance based on experimental results on real data (results shown in the next Section).

For filtering, VEF considers as features the variant annotations specified in the INFO field of the VCF file used for training, as well as the quality score. For example, the types and meanings of the annotations contained in the INFO field of version 4.2 VCF files are as follows:

- AC: Integer, allele count in genotypes.
- AF: Float, allele frequency.
- AN: Integer, total number of alleles in called genotypes.
- DP: Integer, approximate read depth.
- FS: Float, strand bias estimated using Fisher’s exact test.
- MLEAC: Integer, maximum likelihood expectation (MLE) for the allele counts (not necessarily the same as the AC), for each alternate allele, in the same order as listed.
- MLEAF: Float, maximum likelihood expectation (MLE) for the allele frequency (not necessarily the same as the AC), for each alternate allele, in the same order as listed.
- MQ: Float, root mean square of the mapping quality of reads across all samples.
- QD: Float, variant call confidence normalized by depth of sample reads supporting a variant.
- SOR: Float, strand bias estimated by the symmetric odds ratio test.

However, some features may be excluded for training. Specifically, VEF uses the following heuristic method for feature selection. Features are selected unless i) there are more than 10% of missing values for this feature; or ii) variance of this feature is smaller than 0.1. Random forest cannot deal with missing values, and hence during the training phase, variants that contain missing values for at least one of the selected features are dropped (i.e., those variants are not used for training). The first criterion is therefore selected to avoid excluding many variants for training, and the second criterion excludes features that are not informative for filtering.

Once the features are selected, VEF extracts for each considered variant in the training VCF file the values of the used features for that variant, and generates a label indicating whether the variant is true or not. The latter is computed by checking whether the variant is contained in the gold standard or not. We recommend using the script “hap.py” [Krusche *et al.*, 2018] for this purpose, which is able to deal with complex variants. VEF then uses this information to train the random forest. Similarly to the state-of-the-art filtering tools VQSR and HF, VEF trains on the SNPs and INDELs separately, building a separate pre-trained model for each of them.

Once trained, the model can be directly applied to filter the called variants contained in a VCF file (hereafter denoted as the target VCF file). In particular, for each variant in the target VCF file, each DT that composes the RF will output the estimated class (correct or not) of that variant, by traversing the tree using the value of the variant’s features/annotations (represented in blue in Figure 2). The RF will then compute the arithmetic average of the output of all DTs, and generate the final prediction. VEF uses by default a cut-off value of 0.5, i.e., a variant is retained if its probability of being correct is equal to or above 0.5. As state above, VEF assumes that both the target VCF file and the one used for training were generated by the same pipeline. As such, the target VCF file is expected to contain all features used to train the model. Extra features in the target VCF file are discarded, and VEF fails to work when there are features missing. Finally, since RF cannot deal with empty values, features with no value for a given variant are replaced with the average value of that feature in the training dataset, which is stored together with the trained model [Breiman and Cutler, 2004].

The output of VEF is a VCF file that includes a “VEF” score in the INFO field of each variant. This score corresponds to the probability of the variant being correct, calculated by the RF. VEF also specifies ‘VEF_FILTERED’ in the FILTER field of the variants that VEF estimates are incorrectly called. Figure 3 shows the described workflow.

**Figure 3:**
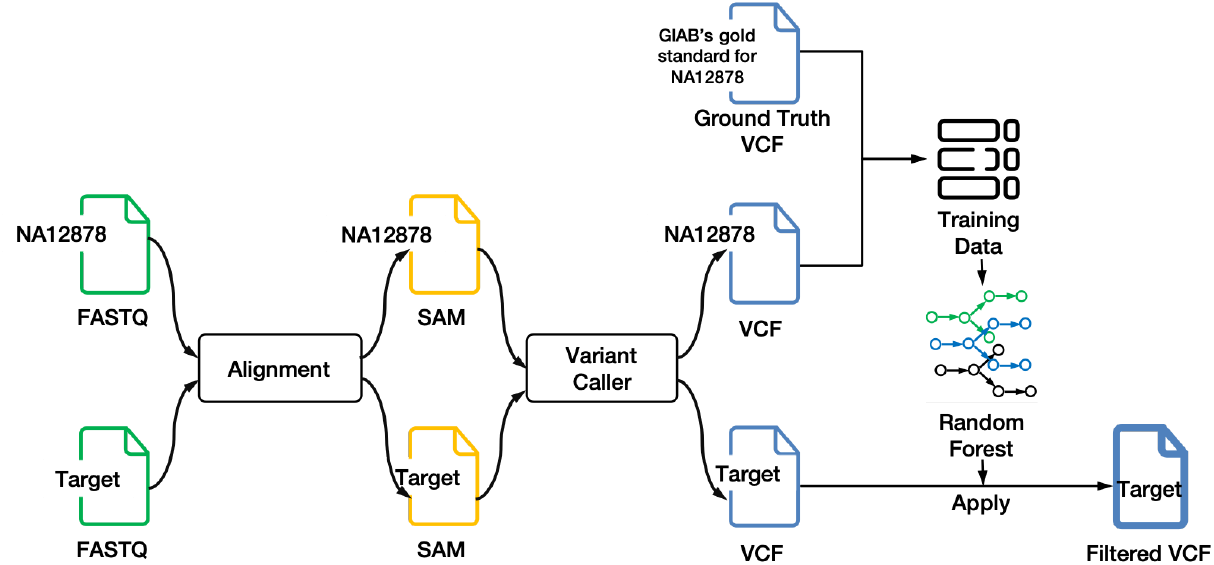
Workflow of VEF for the case where a VCF file from Human sample *NA12878* is used for training. The alignment and variant caller tools used to generate the training and target VCF files are assumed to be the same.

Finally, it should be mentioned that VEF has been designed and optimized to filter variants from single non-cancerous samples, and has only been tested in this setting. As such, other filtering methods may be preferred to filter variants from large cohorts (e.g., VQSR) and/or from cancerous samples.

## 3 Results

In what follows, we first compare the classification accuracy of the different considered ensemble methods, to support the choice of random forest as the filtering method in VEF. We then compare the performance of VEF to that of state-of-the-art filtering tools VQSR and HF. Finally, we provide some additional results to show the robustness of VEF against missing values in the features used for classification, and possible differences in the training and target VCF files with regards to the coverage and the sequencing machine used to generate the raw data.

All experiments were run on a server with Intel(R) Xeon(R) CPU E5-2698 v4 @ 2.20GHz and 515GB of RAM.

### 3.1 Datasets

For the analysis, we only consider datasets for which the *ground truth* containing the set of high-confident variants is available. Note that a ground truth is needed for the training of VEF and the considered ensemble methods, as well as for analyzing the classification accuracy of the considered filtering methods when applied to the target VCF files.

Specifically, we consider data from Human samples *NA12878 (HG001)* and *NA24385 (HG002)*. For these samples (genomes), the NIST’s GIAB consortium has provided the corresponding set of high-confidence variants (*gold standard*), which we use as ground truth [Zook *et al.*, 2018, 2014]. It should be noticed that these golden standards only contain variants located in high-confident regions. The particular datasets used for our study are introduced as needed in the following subsections, together with any possible preprocessing steps applied to them.

### 3.2 VEF: model and parameter selection

To assess the classification accuracy of the considered ensemble methods RF, GBDT, and AdaBoost for inclusion in VEF, we considered evaluation metrics that take into account both the retained variants and the filtered ones, as we are interested in evaluating the performance of binary classifiers. In particular, we consider:

- True positives (TPs): set of *true* variants that were retained;
- False Positives (FPs): set of *false* variants that were retained;
- False Negatives (FNs): set of *true* variants that were filtered out;
- True Negatives (TNs): set of *false* variants that were filtered out.

*True* (*false*) variants are defined as those (not) contained in the gold standard variant set. We are therefore interested in binary classifiers that maximize the number of TPs and TNs while minimizing the number of FPs and FNs.

We further consider the sensitivity (also denoted as recall or True Positive Rate (TPR)) and precision (also denoted as Positive Predictive Value (PPV)) to evaluate the performance of the binary classifiers. They are defined as 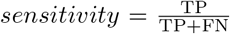 and 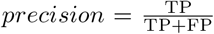. In order to have a single metric, we also consider the *F*_1_ score, given by the harmonic mean of the sensitivity and the precision, i.e., 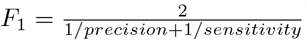. A model with large sensitivity, precision, and *F*_1_ score is desired. When comparing different models, we consider the model with the highest *F*_1_ score to be the best performing one. Unless otherwise stated, these metrics are computed for default cut-off value 0.5.

Nevertheless, ensemble methods can operate at different cut-off values, which may lead to different performance metrics, and thus different operation points. A standard method of evaluation is therefore to compute the ROC curve, obtained by plotting the TPR (defined above) as a function of the False Positive Rate (FPR), defined as 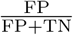, for different cut-off values. Methods with higher Area Under the Curve (AUC) are considered to be better performing.

For the analysis, we used dataset *ERR262997* from Human sample NA12878. This dataset has been generated with an Illumina HiSeq 2000, has 30× coverage, and can be accessed at www.ebi.ac.uk/ena/data/view/ERA207860. We generated the corresponding VCF file from the provided FASTQ files by running GATK’s best practice pipeline (version 3.8.0) [Van der Auwera *et al.*, 2013] with reference genome GRCh37/b37 from the 1000 Genomes Project. Since the considered gold standards only contain variants from the high-confident regions, only variants in these regions are used for training of the ensemble methods.

We first consider the variants found in chromosome 11. In particular, we randomly split the variants into training and testing sets, with an 80%/20% ratio. The main hyper-parameter of the analyzed ensemble methods is the number of base learners (single classifiers), and we first consider 150 and 200 for the study.

Figure 4a summarizes the filtering accuracy of RF, GBDT, and AdaBoost when applied to SNPs. In particular, the figure shows the ROC curve of each classifier. As it may be observed, random forest offers better classification accuracy than GBDT and AdaBoost, since the ROC curves of RF are almost always on the top left of the others, even when using a smaller number of base learners. This is also reflected by the AUCs obtained by each method, with RF having an AUC of 0.943, GBDT 0.940 at most, and AdaBoost 0.934. In addition, the *F*_1_ score of RF, computed for default cut-off value 0.5, is larger than that obtained with GBDT and AdaBoost (for both 150 and 200 learners). Similar results are obtained when considering INDELs instead, as shown in Figure 4b. In particular, RF achieves an AUC of more than 0.922, as compared to an AUC of at most 0.916 for GBDT and 0.904 for AdaBoost. Regarding the *F*_1_ score, RF also exhibits the highest value, with 0.835 for both 150 and 200 learners.

**Figure 4:**
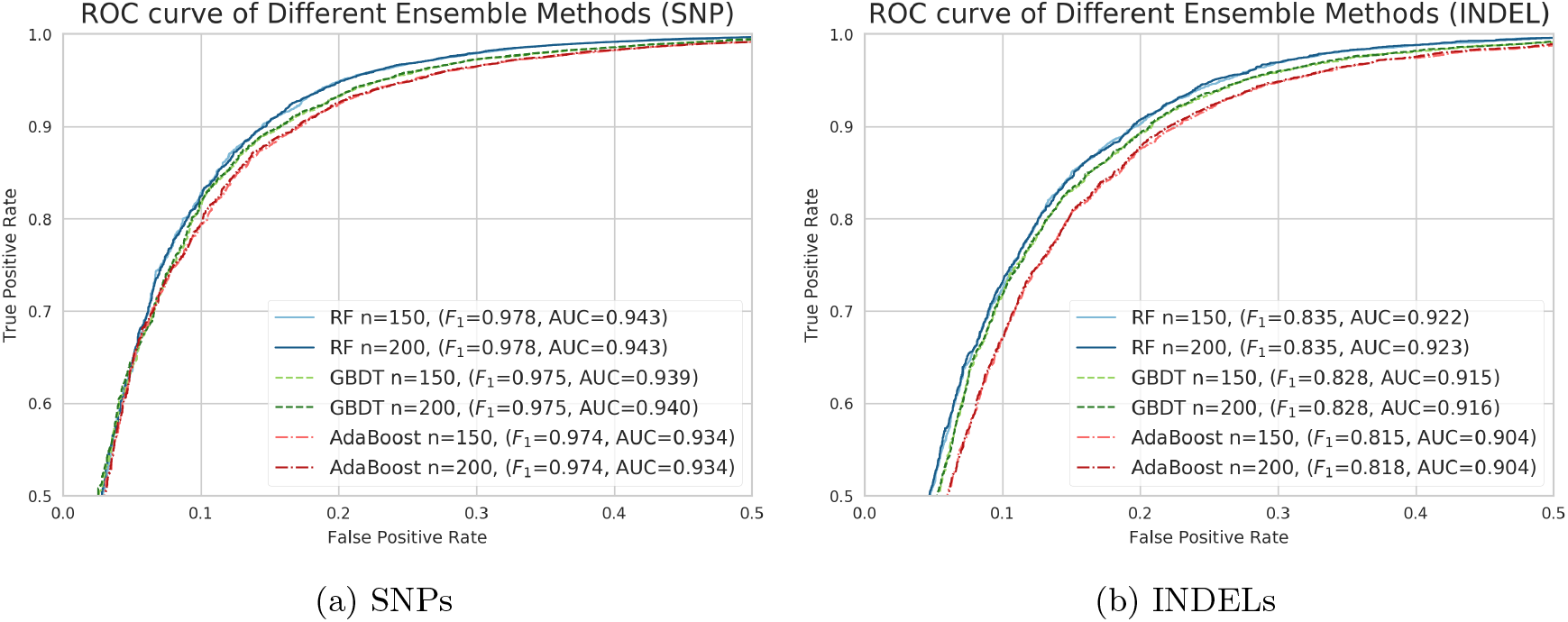
Comparison of ROC curves and AUC values of Random Forest (RF), Gradient Boosting Decision Tree (GBDT) and AdaBoost, when applied to the variants found in chromosome 11 of the ERR262997 dataset. The variants are randomly split into training and testing sets, with a 80%/20% ratio. *n* is the number of single classifiers. Coordinates are tailored to 0.5. The *F*_1_ score is computed for default cut-off value 0.5.

Based on the above results, RF offers the best classification performance, and hence we further tested which number of base learners exhibits the highest *F*_1_ score for this method, when applied to the same dataset. For the analysis, we performed 5-fold cross validation for each of the tested number of base learners, i.e., we first divided the dataset into 5 subsets, and then trained the model on four subsets and validated on the remaining one, iterating over the five subsets. The final score is then computed as the average score over the five rounds. The results for SNPs and INDELs are shown in Figure 5. As can be observed, the performance is consistent among different number of base learners in the RF. In addition, for the SNPs, the fluctuation of the *F*_1_ score mean is less than the standard deviation, which indicates that it is equally good to choose any setting.

**Figure 5:**
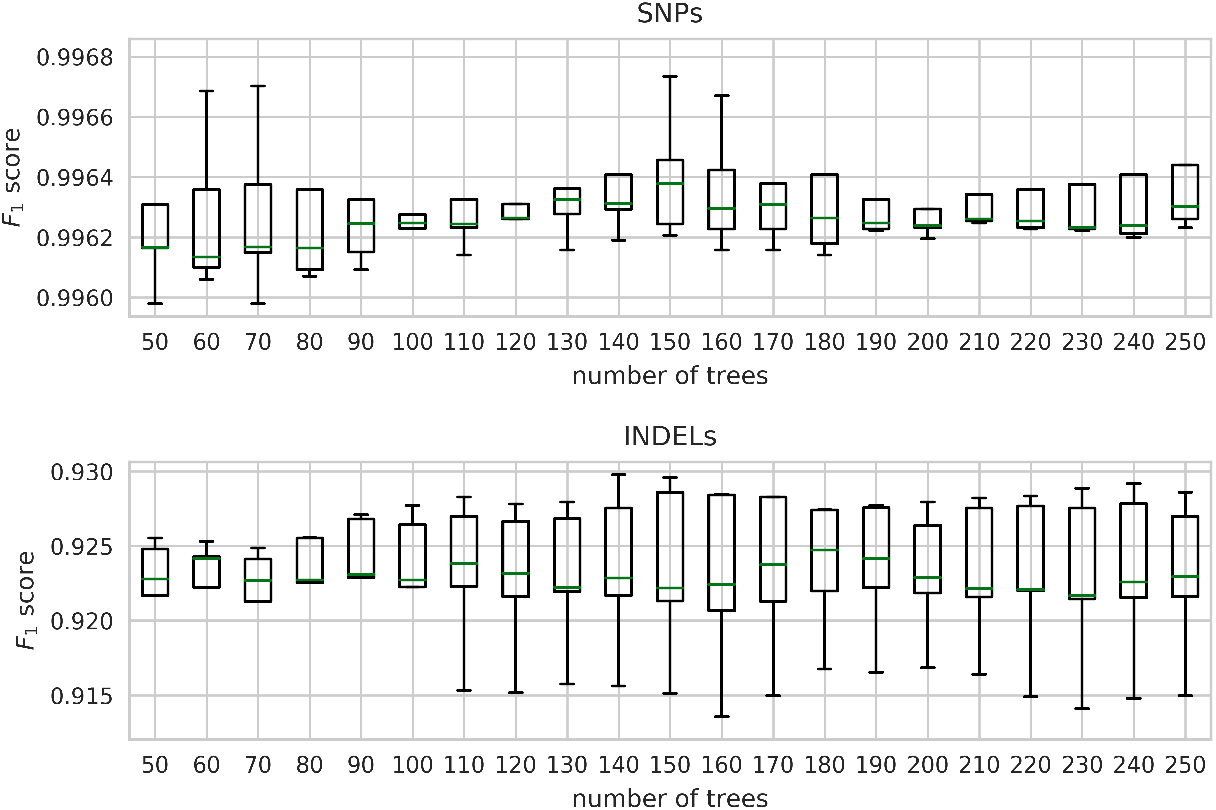
Boxplot (representing median and quartiles) of the *F*_1_ score of RF for different number of base learners when applied to SNPs and INDELs found in chromosome 11 of dataset ERR262997. 5-fold cross validation was used during training. The *F*_1_ score is computed for cut-off value 0.5.

To further validate the higher classification accuracy of RF, we performed another set of experiments. In particular, we trained using the variants from chromosome 11, and tested on the variants from chromosome 20, for the same dataset *ERR262997*. This set of experiments also serves to evaluate the generalizing ability of each method. The resulting ROC curves for each method are depicted in Figures 6a (SNPs) and 6b (INDELs). As can be observed, in both cases RF outperforms GBDT and AdaBoost in classification accuracy. In particular, RF with 150 and 200 base learners achieve an AUC value of 0.963 for SNPs and 0.919 and 0.920 for INDELs, respectively, which are larger than those of GBDT and AdaBoost in both cases. In terms of *F*_1_ score, RF also exhibits the highest value. Regarding the number of base learners for RF, 150 exhibits the same or better classification performance than 200 learners.

**Figure 6:**
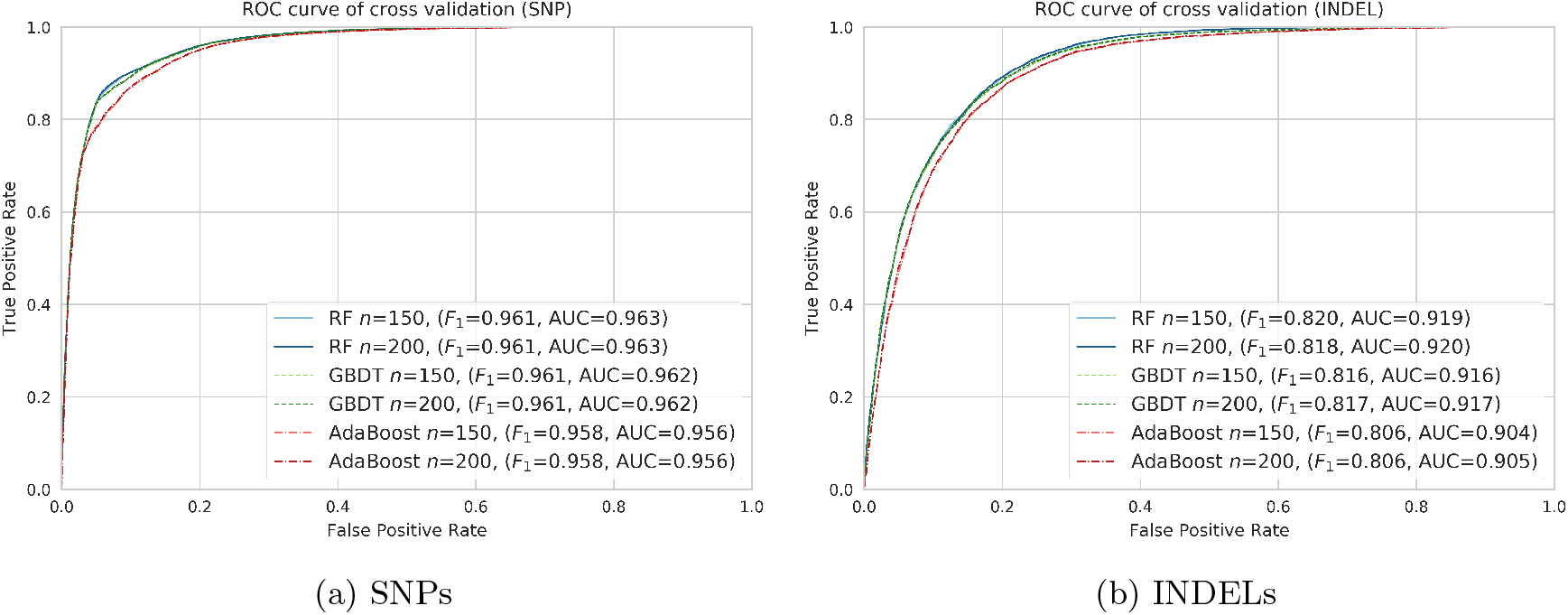
Comparison of ROC curves and AUC values of RF, GBDT and AdaBoost when trained on the variants from chromosome 11 and tested on the variants from chromosome 20, for dataset ERR262997. *n* is the number of single classifiers. Coordinates are tailored to 0.5. The *F*_1_ score is computed for default cut-off value 0.5.

Given these results, in all experiments presented in this paper, VEF is based on RF with a default number of base learners set to 150, and a cut-off value of 0.5. When using the VEF package, however, the number of base learners can be set to a different value, which can be chosen by applying cross-validation to the data, for example.

To further check the classification ability of VEF, we extended the previous experiment by training VEF on the SNPs of each chromosome and testing on the remaining ones. The *F*_1_ score obtained for each case is shown in Figure 7. Diagonal entries are omitted since we are only considering training and testing on different chromosomes. We observe that the choice of chromosome for training has almost no effect on the classification performance of VEF. Finally, we observe that chromosomes 11, 19 and *X* exhibit the lowest *F*_1_ scores, which indicate that the SNPs are more difficult to be classified correctly in these cases.

**Figure 7:**
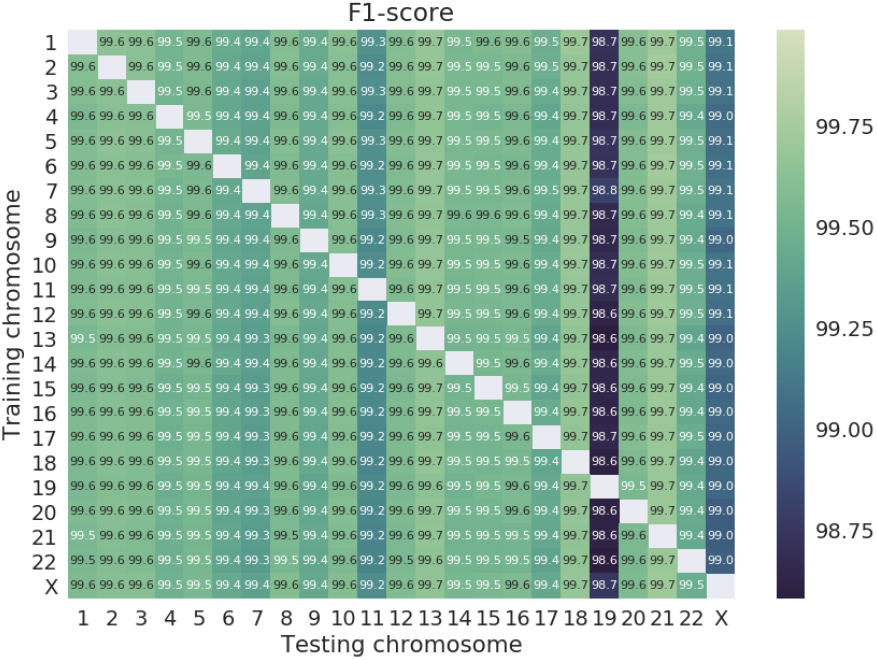
Heatmap of the *F*_1_ score obtained by VEF when trained on the SNPs of a given chromosome and tested on the remaining ones, for all chromosomes of dataset ERR262997. Entry (*i, j*) corresponds to the *F*_1_ score when trained on chromosome *i* and tested on chromosome *j*.

### 3.3 Performance analysis of VEF

After model and parameter selection for VEF, we performed validation on more datasets to test its generalization ability, and how its classification performance compares to that of the state-of-the-art filtering tools VQSR and HF. In all experiments, VQSR and HF are applied following GATK’s recommendations [Broad Institute, 2017, 2018]. Specifically, VQSR is run with threshold 99.5% and the following annotations: DP, QD, SOR, FS, MQRankSum, and ReadPosRankSum. For the HF, the applied cut-offs are “QD < 2.0 or MQRankSum < −12.5 or ReadPosRankSum < −8.0 or SOR > 3.0” for SNPs, and “QD < 2.0 || FS > 200.0 || ReadPosRankSum < −20.0” for INDELs. Finally, the default cut-off of VEF is set to 0.5. When generating the ROC curves, however, VQSR and VEF are run with different thresholds and cut-off values, respectively.

First, we compared the performance of VEF to that of VQSR and HF when applied to chromosome 20 of NA12878 dataset ERR262997. VEF was trained on the variants of chromosome 11. The classification performance of each method is shown in Figures 8a (SNPs) and 8b (INDELs). As can be observed, VEF exhibits improved classification ability over VQSR in this case, for both SNPs and INDELs. This is reflected in the AUC values, 0.963 versus 0.928 for SNPs, and 0.919 versus 0.835 for INDELs. When run with default cut-off values, VEF also exhibits improved performance with respect to VQSR, reflected by a higher *F*_1_ score (0.962 versus 0.938 for SNPs and 0.820 versus 0.803 for INDELs). Nevertheless, for SNPs, the HF achieves the highest *F*_1_ score, given by 0.986. For INDELs, however, the HF achieves the lowest *F*_1_ score.

**Figure 8:**
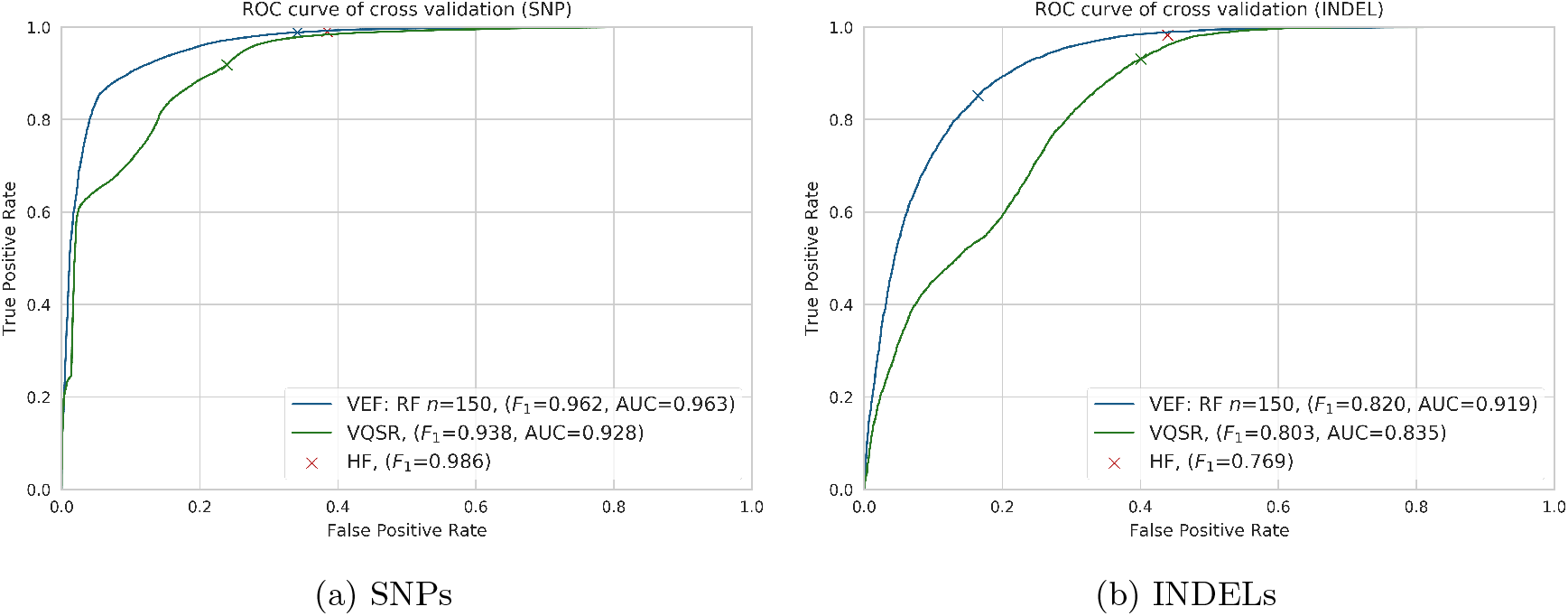
ROC curves of VEF and VQSR when applied to variants from chromosome 20 of the ERR262997 dataset. VEF is trained on chromosome 11. Crosses represent the performance of each method for default cut-off values. Performance of HF is also shown for comparison.

Next, we used WGS aligned data (in the form of BAM files) of Human samples *NA12878* (HG001) and *NA24385* (HG002). The considered datasets were prepared using 10× Genomics’ GemCode and were sequenced with an Illumina HiSeq 2500 machine, and they can be downloaded at ftp://ftp-trace.ncbi.nlm.nih.gov/giab/ftp/data/, followed by NA12878/10XGenomics/ and AshkenazimTrio/HG002_NA24385_son/10XGenomics/, respectively. The coverage is 34× for NA12878 and 25× for NA24385. GATK’s best practice pipeline (version 3.8.0) was used to process the BAM files, discover variants, and generate the final VCF files. We used reference genome *GRCh37/hg19*, since it was the one used to generate the considered BAM files. However, the provided gold standards use reference genome *GRCh37/b37*. Hence, we lifted over the generated VCF files from reference genome hg19 to b37 using the Picard tool *LiftoverVcf*.

Since the considered VCF files are generated with the same pipeline, we expect the numeral features in both files to exhibit similar characteristics. This is indeed the case for most features, as shown in Table 1. Note however that the mean and standard deviation for annotations “QUAL” (quality score) and “DP” (approximate read depth) differ in the two datasets. Nevertheless, in practice, using them increases the generalization ability of VEF.

**Table 1:**
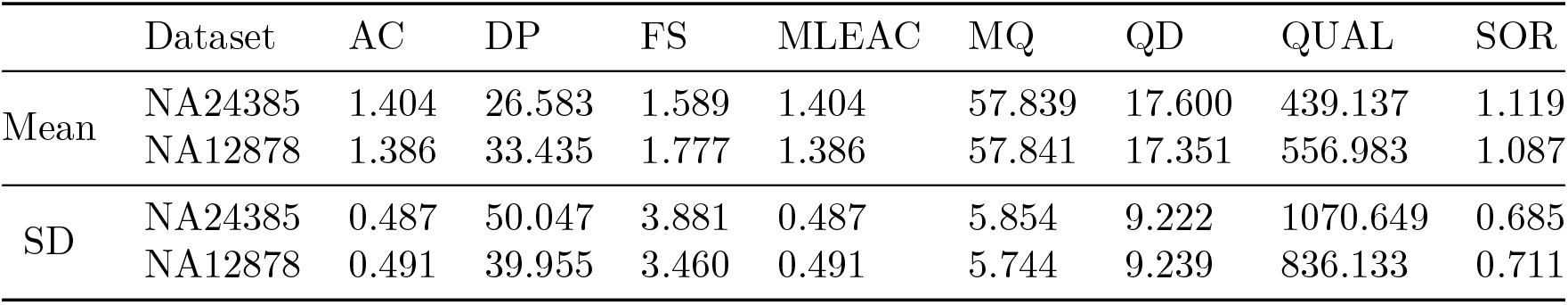
Mean and standard deviation (denoted by SD) of the numerical annotations of the VCF files generated from 10× Genomics WGS datasets NA12878 and NA24385.

|  | Dataset | AC | DP | FS | MLEAC | MQ | QD | QUAL | SOR |
| --- | --- | --- | --- | --- | --- | --- | --- | --- | --- |
| Mean | NA24385 | 1.404 | 26.583 | 1.589 | 1.404 | 57.839 | 17.600 | 439.137 | 1.119 |
|  | NA12878 | 1.386 | 33.435 | 1.777 | 1.386 | 57.841 | 17.351 | 556.983 | 1.087 |
| SD | NA24385 | 0.487 | 50.047 | 3.881 | 0.487 | 5.854 | 9.222 | 1070.649 | 0.685 |
|  | NA12878 | 0.491 | 39.955 | 3.460 | 0.491 | 5.744 | 9.239 | 836.133 | 0.711 |

For the analysis, VEF was first trained using NA12878’s VCF file, and then tested on NA24385’s VCF file, whereas VQSR and HF were directly applied to NA24385’s VCF file. SNPs and INDELs were considered separately. The classification accuracy of VEF, VQSR and HF are shown in Figures 9a (SNPs) and 9b (INDELs). The metrics used are the same as those defined in the previous subsection. In this case, VEF also exhibits better classification accuracy than VQSR, with an AUC of 0.967 as opposed to 0.853 for SNPs, and an AUC of 0.897 instead of 0.625 for INDELs. Note that the improvement in performance is more pronounced for INDELs than for SNPs. In terms of *F*_1_ score, VEF achieves the highest value as compared to VQSR and HF, for both SNPs and INDELs.

**Figure 9:**
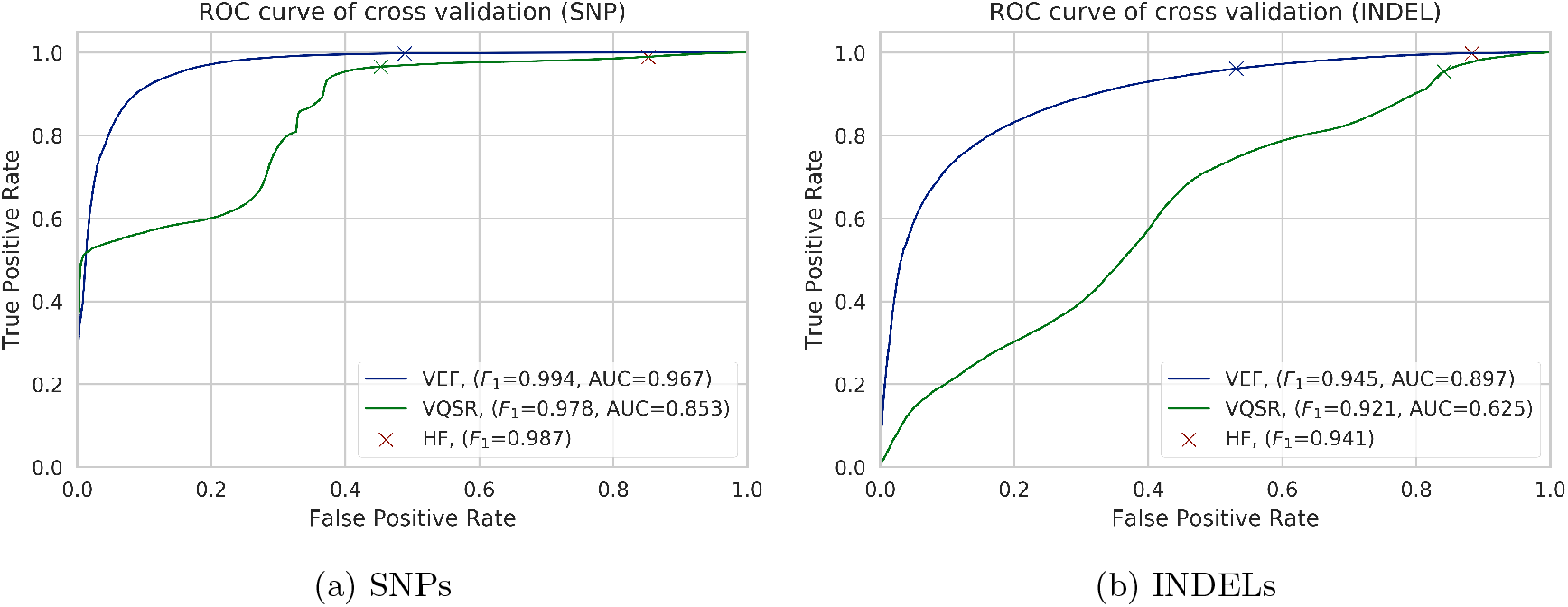
ROC curves of VEF and VQSR when applied to data from WGS Human sample NA24385. VEF is trained on WGS data from Human sample NA12878. Crosses represent classification performance of each method when run with default cut-off values. Performance of HF is also shown for comparison.

The above results showcase the classification ability of each method by analyzing the accuracy of the retained and filtered variants. In the following set of results, however, we directly analyze the accuracy of the filtered VCF files (i.e., the retained variants), by comparing them with the gold standard. We use the benchmark tool “hap.py” [Krusche *et al.*, 2018] for this task. The definition of the metrics computed by this tool differ from the previous ones, as in this case the filtered variants are not considered. In particular, the definition for TPs, FPs, and FNs are as follows.

- True positives (TPs): set of variants contained in both the VCF file and the gold standard;
- False Positives (FPs): set of variants contained in the VCF file but not in the gold standard;
- False Negatives (FNs): set of variants contained in the gold standard but not in the VCF file;

Note that the main difference comes in the definition of the FNs, and the lack of true negatives, as in this case those would correspond to all positions in the genome that do not contain a variant and that are not called in the VCF file. Given the number of TPs, FPs, and FNs, the computation of the sensitivity, precision and *F*_1_ score remains unchanged.

Table 2 shows the sensitivity, precision, and *F*_1_ score for the filtered VCF files generated in the above experiment, as well as for the original VCF file before filtering, for both SNPs and INDELs. Regarding the SNPs, we observe that the VCF file generated by VEF not only achieves the highest *F*_1_ score among the tested filtering tools, but it is also the only filtering method to obtain an *F*_1_ score larger than that obtained with the original VCF file (i.e., the one before filtering). Note, however, that all considered filtering methods improve the precision with respect to the unfiltered VCF file. With respect to INDELs, we observe that in this case both the HF and VEF improve in *F*_1_ score with respect to the unfiltered VCF file, with the HF obtaining the highest value. Similarly to what is observed for the SNPs, all filtering methods improve upon the precision, although there is a decreased in the sensitivity for all methods.

**Table 2:** Comparison of filtering performance of VEF, HF and VQSR when applied to variants from WGS Human sample NA24385. VEF is trained on variants from WGS Human sample NA12878.

| Type | Filter | Sensitivity | Precision | $F_1$ score |
| --- | --- | --- | --- | --- |
| SNPs | Original | 91.3679% | 97.0792% | 94.1370% |
|  | VEF | 91.1771% | 98.0167% | 94.4733% |
|  | HF | 90.4132% | 97.3422% | 93.7498% |
|  | VQSR | 88.3912% | 98.1940% | 93.0351% |
| INDELs | Original | 64.0314% | 82.4307% | 72.0753% |
|  | VEF | 63.7459% | 83.5660% | 72.3226% |
|  | HF | 63.8905% | 83.5940% | 72.4261% |
|  | VQSR | 61.4058% | 84.1312% | 70.9942% |

Finally, Table 3 summarizes the running time employed by VQSR and VEF for filtering the SNPs belonging to WGS Human sample NA24385. The running time of VQSR includes the time needed to run ‘VariantRecalibrator’ (specified as *training*) and ‘ApplyRecalibration’ (specified as *filtering*). VEF was trained on the SNPs from WGS Human sample NA12878. In addition to the improved classification accuracy of VEF with respect to VQSR, VEF also offers significant savings in running time, since once trained, the model can be directly applied to the target VCF files. In this example, the total running time of VQSR is about 49 minutes, whereas once trained, VEF needs less than 4 minutes to perform the filtering. In terms of memory consumption, VEF uses about 5 times less memory than VQSR.

**Table 3:** Running time and memory consumption of VQSR and VEF when applied to the SNPs of WGS Human sample NA24385. VEF is trained on WGS Human sample NA12878.

|  |  | Running Time | Memory |
| --- | --- | --- | --- |
| VQSR | Training | 47 minutes | 12,341 MB |
|  | Filtering | 2 minutes | 10,347 MB |
| VEF | Training | 46 minutes | 2,595 MB |
|  | Filtering | 3.5 minutes | 1,745 MB |

### 3.4 Additional experiments

In order to further analyze the performance of VEF, we conducted three additional tests that focus on dealing with missing feature values, low coverage datasets, and cross sequencing machine usage. We also conducted an additional experiment to analyze the effect of the number of base learners on the classification accuracy of VEF. Due to space constraints, only results for SNPs are shown here. Unless otherwise stated, all tests were conducted using the WGS datasets from Human samples NA12878 and NA24385 used in the previous experiments. All filtering methods are applied to sample NA24385, and VEF is trained on variants from sample NA12878.

#### Missing features

This test is aimed to test the robustness of VEF when some features have missing values for a small subset of the variants. The test was performed by deleting values of features uniformly at random in the target VCF file, with probabilities equal to 5%, 7.5%, and 10%. The python script used for this task is called *kick_out.py* and it is available in the provided GitHub link. VEF, VQSR and HF were then run on the resulting VCF files. For each of the considered probabilities, we generated four VCF files. Table 4 shows the average *F*_1_ score obtained by each of the filtering methods. The hap.py script was used to generate the results. As expected, the *F*_1_ score achieved by VEF decreases as the number of missing values increases. This is not the case, however, for VQSR and HF. Note that one of the main differences between VEF and VQSR is that the latter trains on the target VCF file, which may explain why the performance of VQSR does not deteriorate with the increased percentage of missing values. In fact, VQSR’s performance improves. Nevertheless, VEF obtains the highest *F*_1_ score in all cases, even when compared to the unfiltered VCF file.

**Table 4:**
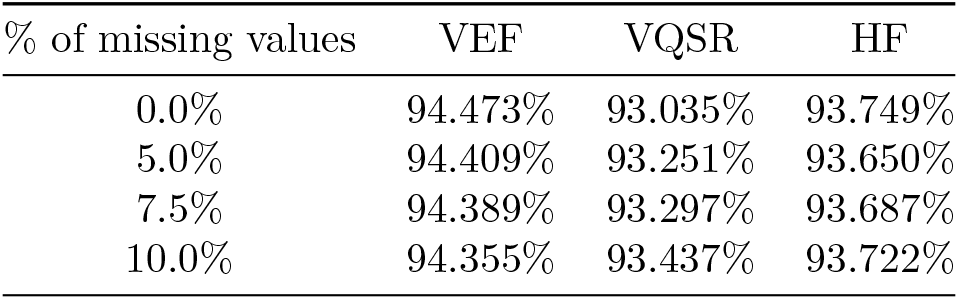
Average *F*_1_ score obtained by VEF, VQSR and HF when applied to the SNPs of Human sample NA24385 with varying percentage of missing features values. The *F*_1_ score of the unfiltered VCF file is 94.137%.

| % of missing values | VEF | VQSR | HF |
| --- | --- | --- | --- |
| 0.0% | 94.473% | 93.035% | 93.749% |
| 5.0% | 94.409% | 93.251% | 93.650% |
| 7.5% | 94.389% | 93.297% | 93.687% |
| 10.0% | 94.355% | 93.437% | 93.722% |

#### Low coverage

In this test we perform filtering on a VCF file generated from low coverage data, serving two aims. First, to evaluate how VEF performs when trained and tested on variant sets generated from data with different coverages, and second, to check the potential of VEF to improve variant accuracy in this case, as low coverage data hinders variant discovery.

For the analysis, we first downsampled the BAM file of sample NA24385, with average coverage 25×. After downsampling, the resulting average coverage is 8.3×. The GATK’s best practice pipeline was then applied to the downsampled BAM file to generate the target VCF file used for filtering. VEF was trained on the VCF from WGS Human sample NA12878, with average coverage 34×. The *F*_1_ scores obtained by VEF, VQSR and HF, as well as the unfiltered VCF file, are summarized in Table 5. We observe that VEF is the only filtering method that improves the *F*_1_ score of the resulting VCF file with respect to the unfiltered one. This result also suggests that VEF performs well even when trained on and applied to variant sets generated from samples with different coverages.

**Table 5:** *F*_1_ score obtained by VEF, VQSR and HF when applied to the SNPs of downsampled WGS Human sample NA24385 (coverage 8.3×). VEF is trained on the SNPs from WGS Human sample NA12878 with coverage 34×.

| Unfiltered | VEF | VQSR | HF |
| --- | --- | --- | --- |
| 72.541% | 72.984% | 71.692% | 72.102% |

To further validate this claim, we performed another test in which VEF is trained on a dataset of lower coverage than that of the target VCF file. For the analysis, we downsampled the BAM file of WGS Human sample NA12878, with coverage 34×, resulting in an average coverage of 10.6×. The resulting BAM file was then used to generate the VCF file in which VEF was trained. All filtering methods were applied to the VCF file generated from WGS Human sample NA24385 with coverage 25×. The results are summarized in Table 6. In this case we also observe that VEF is the only filtering method to improve upon the unfiltered VCF file, increasing the *F*_1_ score from 94.137% to 94.334%. This result supports the claim that VEF does not need to use datasets of very similar coverage for training and testing.

**Table 6:** *F*_1_ score obtained by VEF, VQSR and HF when applied to the SNPs of WGS Human sample NA24385 (coverage 25×). VEF is trained on the SNPs from downsampled WGS Human sample NA12878 (coverage 10.6×).

| Unfiltered | VEF | VQSR | HF |
| --- | --- | --- | --- |
| 94.137% | 94.334% | 93.02% | 93.750% |

#### Cross machine

In this test we evaluate how VEF performs when trained and tested on variant sets generated with the same processing pipeline, but that use raw data produced by different sequencing machines. We used NovaSeq v2 data from the project “NovaSeq S2: Nextera DNA Flex (8 replicates of NA12878)”, which can be found on Illumina’s BaseSpace. We combined data from two lanes, for a total coverage of about 100×. The resulting VCF file was generated using GATK’s best practice pipeline, and it is the file used for training of VEF. VEF was then applied to the SNPs of WGS Human sample NA24385, sequenced by an Illumina HiSeq 2500. The results are shown in Table 7. As it can be observed, VEF achieves the highest *F*_1_ score, demonstrating once more its generalization ability.

**Table 7:** *F*_1_ score obtained by VEF, VQSR and HF when applied to the SNPs of WGS Human sample NA24385, sequenced by Illumina HiSeq 2500. VEF is trained on WGS Human sample NA12878, sequenced by NovaSeq S2.

| Unfiltered | VEF | VQSR | HF |
| --- | --- | --- | --- |
| 99.605% | 99.796% | 99.464% | 99.739% |

#### Number of base learners

To further analyze the effect of the number of base learners in VEF, we performed 5-fold cross validation on the SNPs of WGS Human sample NA12878 sequenced by 10× Genomics. The evaluation is the same as that done for dataset ERR262997, and in fact, similar results are obtained in this case. In particular, we observe (results not shown) that the *F*_1_ score fluctuation is less than the standard deviation, which suggests that the choice of the number of base learners has little effect on the performance of VEF.

## 4 Discussion and Conclusion

With the goal of decreasing the number of incorrectly called variants contained in VCF files, filtering tools such as GATK’s Variant Quality Score Recalibration (VQSR) and Hard Filtering (HF) are applied to the resulting VCF files. To overcome the main drawbacks of these methods, as well as to improve upon them, we have presented VEF, a novel filtering tool based on supervised learning. In particular, VEF trains a Random Forest (RF) on a variant call set from a sample for which a high-confidence set of “true” variants (i.e., a *ground truth* of *gold standard*) is available. Once trained, the model can be applied to filter any VCF file.

The results presented in the previous section demonstrate the improved classification performance obtained by VEF with respect to VQSR and HF, for both SNPs and INDELs. We have also shown that VEF generalizes well, in that it can be trained and applied to VCF files generated from data of different coverages, as well as data produced by different sequencing machines. In addition, VEF has shown to be robust against missing values of features. Another advantage of VEF is that once trained, it can be applied to VCF files of arbitrary size, and no input is needed from the user. VEF also offers significant improvement in running time as compared to VQSR, since once trained, it can be directly applied to filter the variants contained on the target VCF files.

As an added advantage, the use of random forest can provide insights about the significance of each of the features used for training. Specifically, the *gini importance* of a feature is calculated by the average decrease of impurity among all trees for that feature, where the decrease of impurity is computed as the sum of decrease of gini impurity at each splitting node in a tree multiplied by a weight given by the sample ratio at the node [Breiman *et al.*, 1984]. The gini importance is calculated while training. Table 8 shows the gini importance (or Mean Decrease in Impurity) of each feature when VEF is trained on the SNPs of WGS Human sample NA12878. The gini importance of all features sums up to one, and a higher gini importance corresponds to a more important feature. From the provided Table we observe that annotations AC and MLEAC are less important than other features, and hence dropping them or modifying their value would have less impact in the final performance than other annotations (features). Hence, VEF could potentially be used to determine which features are more important. This information could be then used in other applications, such as variant callers, or to decide which features to use for filtering with VQSR and/or HF.

**Table 8:** Gini importance of features (relative score) when VEF is trained on the SNPs of WGS Human sample NA12878.

|  | AC | DP | FS | MLEAC | MQ | QD | QUAL | SOR |
| --- | --- | --- | --- | --- | --- | --- | --- | --- |
| Importance | 0.007 | 0.111 | 0.080 | 0.009 | 0.179 | 0.294 | 0.187 | 0.133 |

Taking all together, VEF could have the potential to replace VQSR and HF as the state-of-the-art filtering tools. It should be noticed, however, that VEF has been designed and optimized to filter variants of single non-cancerous samples. As such, VQSR may be preferred for filtering variants of large cohorts, since it has also been optimized for this setting. Similarly, specialized tools such as Strelka2 [Kim *et al.*, 2018] may be considered for the filtering of somatic variants.

## Acknowledgements

The authors would like to thank Mikel Hernaez for insightful discussions. This work has been partially supported by grant 2018-182799 from the Chan Zuckerberg Initiative DAF, an advised fund SVCF, and a Strategic Research Initiatives (SRI) grant and a CompGen fellowship from UIUC.

## References

Breiman, L. (2001). Random forests. Machine Learning, 45(1), 5–32.

Breiman, L. and Cutler, A. (2004). Random forests - classification description. https://www.stat.berkeley.edu/~breiman/RandomForests/cc_home.htm.

Breiman, L. et al. (1984). Classification and regression trees.

Broad Institute (2017). (howto) apply hard filters to a call set. https://software.broadinstitute.org/gatk/documentation/article.php?id=2806.

Broad Institute (2018). Which training sets / arguments should i use for running vqsr? https://software.broadinstitute.org/gatk/documentation/article.php?id=1259.

Consortium, G. P. et al. (2015). A global reference for human genetic variation. Nature, 526(7571), 68.

Consortium, I. H. et al. (2003). The international hapmap project. Nature, 426(6968), 789.

Danecek, P. et al. (2011). The variant call format and vcftools. Bioinformatics, 27(15), 2156–2158.

DePristo, M. A. et al. (2011). A framework for variation discovery and genotyping using next-generation dna sequencing data. Nature genetics, 43(5), 491–498.

Freund, Y. and Schapire, R. E. (1997). A decision-theoretic generalization of on-line learning and an application to boosting. Journal of computer and system sciences, 55(1), 119–139.

Friedman, J. H. (2001). Greedy function approximation: a gradient boosting machine. Annals of statistics, pages 1189–1232.

James, G. et al. (2013). An introduction to statistical learning, volume 112. Springer.

Kim, S. et al. (2018). Strelka2: fast and accurate calling of germline and somatic variants. Nature methods, 15(8), 591.

Krusche, P. et al. (2018). Best practices for benchmarking germline small variant calls in human genomes. bioRxiv, page 270157.

Li, H. et al. (2009). The sequence alignment/map format and samtools. Bioinformatics, 25(16), 2078–2079.

Ochoa, I. et al. (2016). Effect of lossy compression of quality scores on variant calling. Briefings in bioinformatics, 18(2), 183–194.

Van der Auwera, G. A. et al. (2013). From fastq data to high-confidence variant calls: the genome analysis toolkit best practices pipeline. Current protocols in bioinformatics, 43(1), 11–10.

Zook, J. et al. (2018). Reproducible integration of multiple sequencing datasets to form high-confidence snp, indel, and reference calls for five human genome reference materials. bioRxiv, page 281006.

Zook, J. M. et al. (2014). Integrating human sequence data sets provides a resource of benchmark snp and indel genotype calls. Nature biotechnology, 32(3), 246.

